# Predicting Alzheimer’s Disease Progression from Sparse Multimodal Data by NeuralODE Models

**DOI:** 10.1101/2025.08.26.672412

**Authors:** Andrea Zanin, Stefano Pagani, Mattia Corti, Valeria Crepaldi, Giuseppe Di Fede, Paola F. Antonietti, the Alzheimer’s Disease Neuroimaging Initiative ADNI

## Abstract

Alzheimer’s disease shows significantly variable progressions between patients, making early diagnosis, disease monitoring, and care planning difficult. Existing data-driven Disease Progression Models try to tackle this issue, but they usually require sufficiently large datasets of specific diagnostic modalities, which are rarely available in clinical practice. Here, we introduce a new modeling framework capable of predicting individual disease trajectories from sparse, irregularly sampled, multi-modal clinical data. Our method uses (recurrent) Neural Ordinary Differential Equations to determine the current hidden state of a patient from sparse past exams and to forecast future disease progression, illustrating how biomarkers evolve over time. When applied to the ADNI clinical cohort, the model detected early signs of disease more accurately than common data-driven alternatives and effectively tracked changes in biomarker trajectories that align with established clinical knowledge. This provides a versatile tool for accurate diagnosis and monitoring of neurodegenerative diseases.

## Main

Nowadays, Alzheimer’s Disease (AD) is the leading cause of dementia worldwide [1] and affects approximately 8.45 million people in Europe [2], with a life expectancy of 7.2 years from the start of symptoms[3]. AD is a neurodegenerative disease characterized by predominantly amnestic progressive cognitive decline, resulting from a complex multiscale interplay of genetic, environmental, and lifestyle factors [1]. At the cellular level, the disease features the accumulation of β-amyloid (Aβ) in the form of oligomers, fibrils, and plaques [4], followed by the formation of neurofibrillary tangles made of hyperphosphorylated tau protein [5]. A delay of many years occurs between the beginning of these molecular processes (preclinical stage) and the onset of the first visible symptoms [1]. Moreover, early clinical manifestations are highly varied, and differential diagnosis is difficult due to the overlap with other neurodegenerative disorders causing cognitive decline. Beyond the human cost, AD imposes a significant economic burden: in Europe, the annual cost of AD care is estimated at 141 billion euros [2].

Both the FDA and the EMA have approved a novel disease-modifying treatment (Lecanemab), and a second monoclonal antibody against Aβ (Donanemab), approved by the FDA, recently received a positive recommendation for marketing authorization by the EMA (EMADOC-628903358-125940). However, the appropriate prescription of such disease-modifying treatments requires early and accurate diagnosis based on risk factor assessments and biomarker analysis to support the AD biological substrate [6], which is essential for the identification of candidates to the novel therapies. Additionally, this highlights the clinical need for ongoing monitoring of disease progression, treatment effectiveness, and safety. To address these aspects, diagnosis and treatment of AD rely on a broad spectrum of multimodal data: neuropsychological assessments, neuroimaging, blood tests, genetic data, and cerebrospinal fluid (CSF) analysis [1,2,7]. Ideally, most tests would be conducted regularly to track disease progression, but invasiveness and costs make this not feasible. Moreover, AD biomarkers [8] and symptoms [9] vary significantly among patients. In this context, machine learning (ML) techniques that analyze patient data can improve diagnosis and monitoring. Specifically, data-driven disease progression models (DPMs) are an emerging class of ML tools that predict how the disease will develop over time [10]. They are based on a patient’s exams (for model personalization) conducted up to the last visit and a longitudinal dataset covering multiple known disease pathways. DPMs can help analyze complex patterns in multimodal data for early diagnosis, personalized monitoring, treatment, and guiding clinical trial design.

Several practical and technical challenges arise when developing ML models for neurodegenerative diseases due to the high fragmentation of patient data. The scattered examinations, collected over different timeframes, provide only a partial view of the disease’s longitudinal progression. This limitation is further accentuated by the significant inter-patient variability, particularly in the early stages of the disease, where differential diagnosis is still an open challenge [11]. In this context, a DPM is expected to incorporate multiple scattered biomarkers and cognitive scores, capture disease progression with subtype differentiation (rather than reducing it to a single trajectory) and support applications to personalized medicine.

In this work, we introduce a novel time-continuous DPM that can predict the progression of a wide range of AD markers, aiding neurologists in offering personalized diagnoses to patients and anticipating their clinical trajectories across the disease progression. Our model is based on a new hybrid architecture called ODE-AE, which combines Ordinary Differential Equations (ODEs) and neural networks to form an auto-encoder (AE) structure, allowing it to handle sparse observations of various biomarkers. The separate components forming our hybrid architecture have already been applied to AD. ODE models have been used to model AD disease progression, but they usually consider each biomarker separately or, at most, in small groups (see [10] and references therein); therefore, they do not provide a unified disease timeline and stage for each patient. Prior work has successfully applied neural network models to AD patient classification based on MRI imaging, PET imaging, and neuropsychological assessments [12]. Deep learning models have also been used for AD disease progression modeling in both unimodal [13–16] and multimodal contexts [17]; however, these models can process only a few data modalities and are not designed for irregularly sampled data. In recent years, VAEs have become a robust generative model architecture for multimodal AD DPMs [16,18]. VAEs have been combined with recurrent neural networks [18] or parametric trajectories [16] to incorporate a temporal structure into the model. The longitudinal VAE proposed in [16] is a latent-time regression approach (as defined in [10]): the model jointly learns typical biomarker trajectories along a data-driven latent time axis and the warping functions that map each patient’s timeline to the latent time. Conversely, the MC-RVAE model [18] uses a recurrent neural network to directly model the temporal evolution in the natural time scale of the data. Our novel ODE-AE model combines all these features into a single architecture consisting of an encoder and a decoder. The encoder uses a modified ODE-recurrent neural network (ODE-RNN) to learn the past temporal evolution of a latent probabilistic representation of a patient, from the last examination back to the first. The decoder, in a generative manner, samples from the distribution of the latent representation identified by the encoder, forecasts it with a Neural ODE, and then reconstructs the corresponding dynamics of AD markers with a standard feed-forward neural network (DECODER-NN). A schematic overview is given in Figure 1.

**Figure 1.**
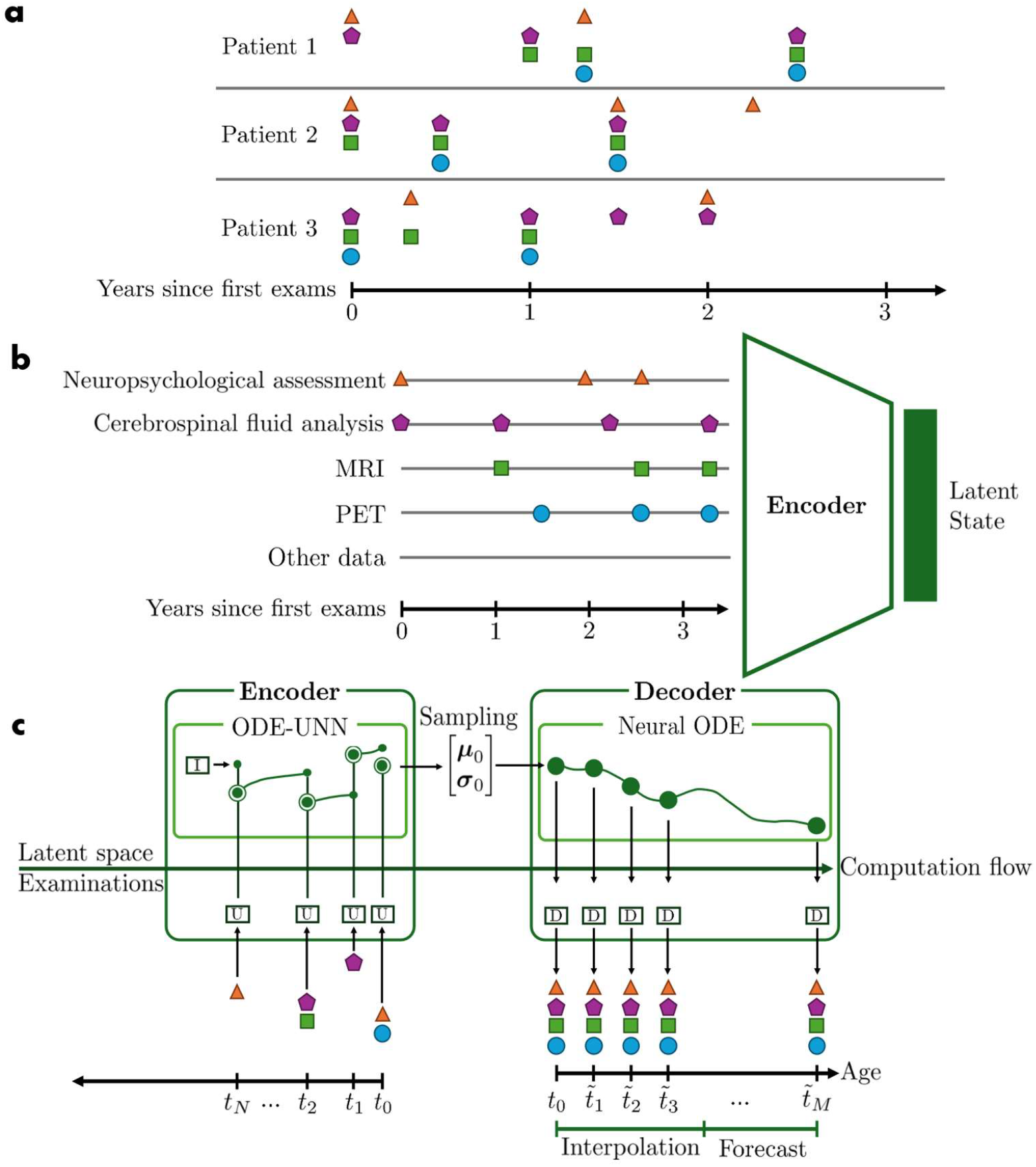
Schematic representation of the NeuralODE-based DPM. **a**, conceptual representation of available scattered data for three different patients, with each symbol representing a distinct category of clinical exams (Neuropsychological assessment, cerebrospinal fluid analysis, MRI, PET, other data). **b**, ENCODER architecture that compresses sparse time series into a dense latent state, encoding the patient’s health status. **c**, ODE-AE architecture that predicts exam results at each time point. I, U, and D stand for INIT-NN, UPDATE-NN, and DECODER-NN, respectively.

For any given patient, each visit, including baseline and follow-up assessments (i.e., the multimodal panel of examinations), is represented by a vector **x**_*i*_ ∈ (ℝ ∪ {NA})^*K*^. Our model represents the patient state at time *t* as a latent vector **z**(*t*) ∈ ℝ^*d*^. To identify a suitable distribution of initial conditions (i.e., the latent state of the patient at the first visit), the variational encoder architecture employs a hybrid architecture based on an ODE-RNN [19]. Here, the RNN is modified to allow for both compressing the visit data **x**_*i*_ into the latent manifold and updating the current latent state with the visit information (we refer to it as UPDATE-NN). The neural ODE [20] part instead fills the gap within two consecutive visits by evolving in time the latent state. The output of the encoder **Z** ∈ ℝ^2*d*^ is interpreted as the concatenated mean ***μ***_0_ and the diagonal covariance ***σ***_0_ of a multivariate Gaussian distribution, which makes the model generative. From a sample of the latter, the latent state is evolved over time integrating a NeuralODE, and an additional fully connected neural network finally maps *z*(*t*) into **x**(*t*) that is the prediction of the visit vector from the latent state. The combination of these two architectures forms the decoder. The complete model architecture, represented in Figure 1, is formalized as:

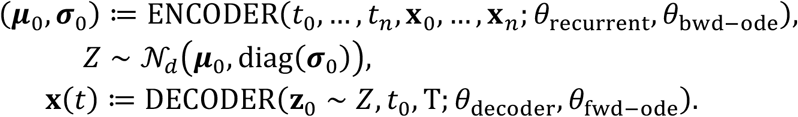

Here *θ* = {*θ*_recurrent_, *θ*_wd–ode_, *θ*_decoder_, *θ*_fwd–ode_} are the parameters (weights and biases) of our architecture that are trained simultaneously in an end-to-end framework. Thanks to its unique design, which merges ODEs and neural networks, our model can handle input data that was sampled irregularly, has high percentages of missing values, and even misses some features entirely, which makes it suitable for clinical applications. The model is trained in an unsupervised fashion with masked loss function components (see Methods), which allows us to fully take advantage of the available clinical data. The model simultaneously predicts how a broad range of AD markers evolve, allowing it to capture complex interactions among multiple markers. We implemented several techniques to investigate the latent patterns learned by our model, which can help interpret the results and guide future research on AD pathology.

Additionally, the model’s continuous output enables it to represent a broad spectrum of neurodegenerative patterns rather than just a few discrete diagnoses, providing a personalized and quantitative prognosis for the patient. We designed the model to be highly flexible and adaptable to various tasks and different types of dementia, as well as other neurodegenerative diseases. It can be used as a regression model to forecast disease progression, estimate results for tests not yet performed, and assist in diagnosis. We also constructed a classification model for patients’ stratification by equipping the ENCODER and the Neural ODE architectures with an additional feed-forward neural network with SoftMax output activations. In summary, our model offers a unique combination of handling highly multimodal, sparse, and irregular data, continuous disease modeling, and flexible prediction capabilities—features not found together in any other model in the literature.

## Results

In this section, we report the results obtained with the proposed model using data from the Alzheimer’s Disease Neuroimaging Initiative (ADNI) database (adni.loni.usc.edu). Specifically, we constructed a multimodal dataset of 46 different features extracted from the ADNI1, ADNI GO, ADNI2, and ADNI3 cohorts, which followed different exam compositions and schedules. A short description of the ADNI architecture is provided below and further detailed in the Methods section. The Alzheimer’s Disease Neuroimaging Initiative (ADNI) dataset is organized to support the longitudinal study of multimodal biomarkers on the entire spectrum of Alzheimer’s disease. Data is indexed by assigning each participant a unique identifier (RID) and Patient ID (PTID), where PTID encodes site and subject information. Longitudinal visits are tracked using standardized visit codes (VISCODE and VISCODE2) such as bl (baseline), m12 (12 months), or SC (screening), accompanied by actual acquisition dates. Subject characteristic, neuroimaging data, clinical and cognitive assessments, biofluid biomarkers and genetic and other-omics have been collected and stored following the relative clinical protocols. A summary of the ADNI data structure and the list of features extracted for the training of our model are reported in Table 1.

**Table 1.**
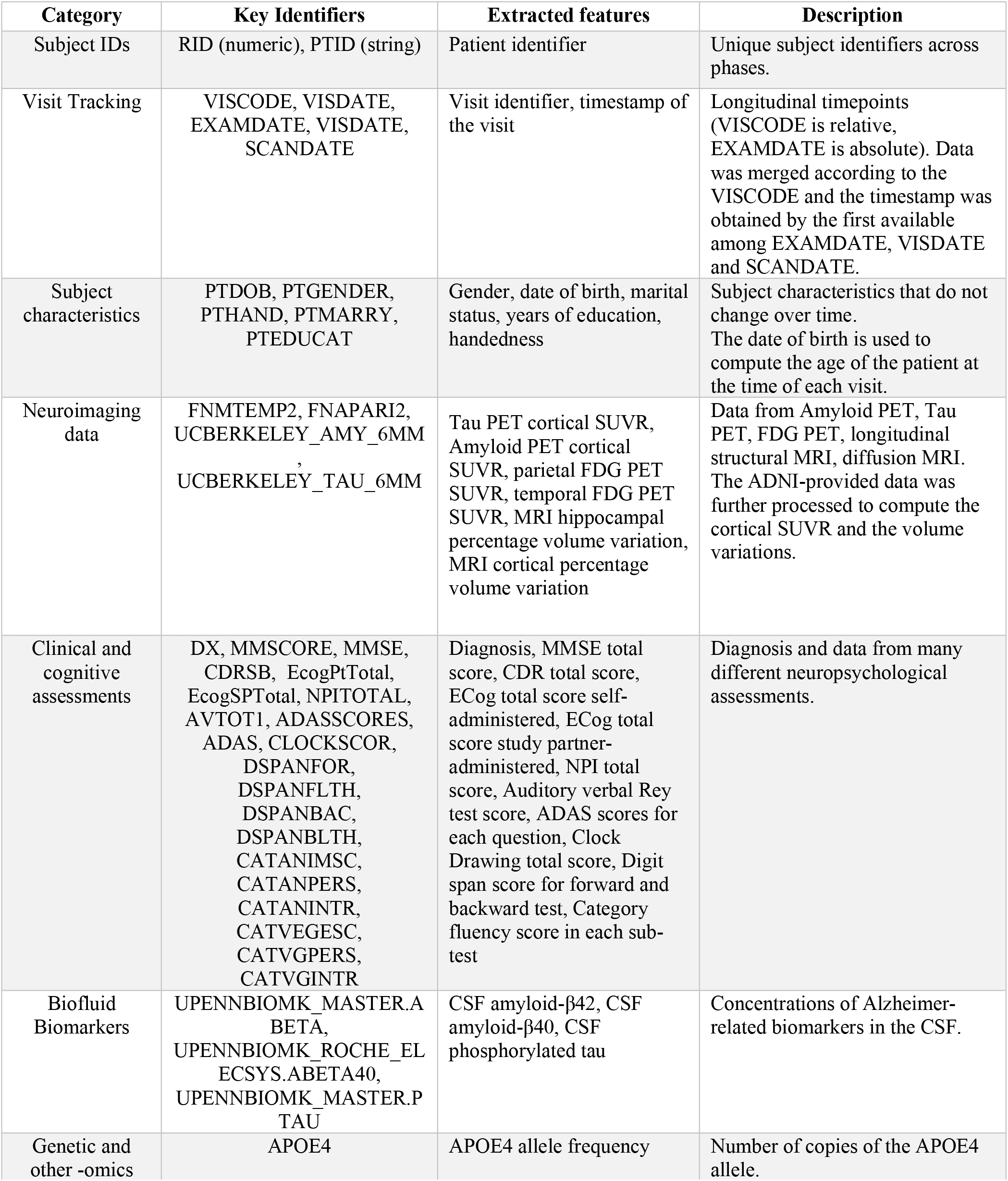
Overview of ADNI data structure.

### Early diagnosis

A crucial task for clinicians is early diagnosis, which can be defined as predicting whether a patient who, using the standard criteria, is currently diagnosed as healthy, will remain healthy or develop some form of cognitive impairment in the future. An early diagnosis allows choosing a proper early intervention, which can delay or even prevent the onset of dementia [21]. In our test set, 310 out of the 948 currently healthy patients (33%) change diagnosis at some point in the future, so the number of patients that could benefit from better early diagnosis techniques is large. Our classification model has been designed to subdivide patients into the healthy, MCI, and dementia groups at each time step. This allows us to determine the patient’s current health status and predict if and when it will change based on the predicted latent trajectory. To train the classification model, we loaded the weights from the complete model and initialized the new decoder weights, then trained the decoder until convergence, while keeping the other weights fixed. Finally, we trained the entire network end-to-end. We trained the classification model using the truncated time series of all the patient data up to the present as input (excluding the diagnosis), and the whole time series of the diagnoses as output. We used the following loss function:

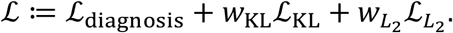

Here *ℒ*_diagosis_ is the sum of the categorical cross-entropy between the predicted and the proper diagnosis at each time step, where the appropriate diagnosis is available; In contrast, the KL and the *L*_2_ loss are defined in the ‘Training’ section (see Methods). Using our model on currently healthy patients, we can predict whether they will develop some form of cognitive impairment with 74% precision, 52% recall and 79% accuracy (the associated confusion matrix is reported in Figure 2a). Finally, we have tested two other approaches to predict the patient diagnosis: the diagnosis predicted by the original model, and the predicted MMSE or CDR scores to choose a diagnosis according to fixed cut-offs. All these approaches have significantly lower accuracy than the fine-tuned classification model: 69% for the original model, 73% for the MMSE-based method and 74% for the CDR-based method.

**Figure 2.**
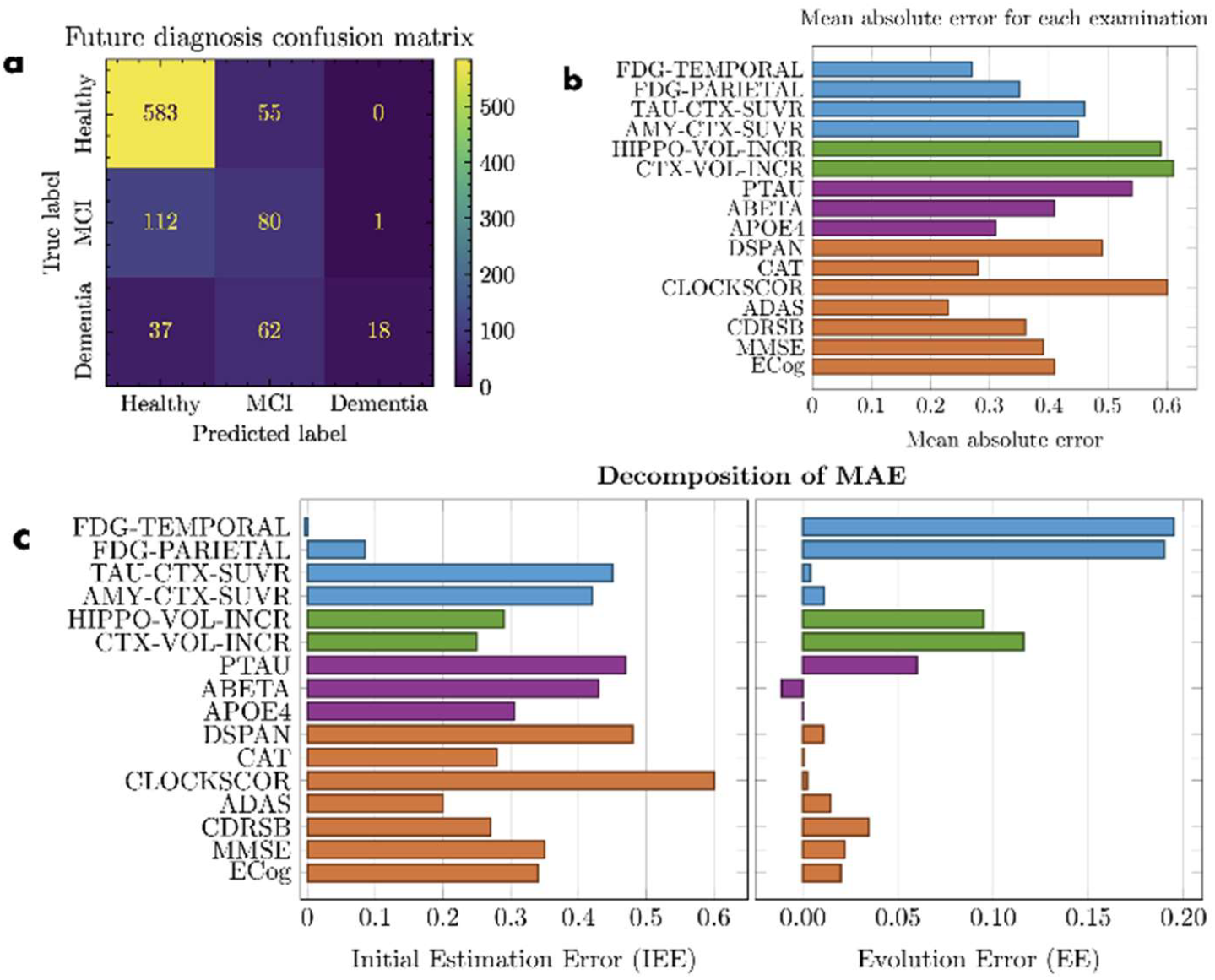
Early diagnosis and feature reconstruction analyses of the model. **a**, confusion matrix for the future diagnosis of currently healthy patients. **b**, mean absolute error for the reconstruction of several examinations, computed on normalized features. **c**, decomposition of mean absolute error into initial estimation error and evolution error. The list of acronyms used is detailed in Table 4.

### Biomarker reconstruction analysis

To understand the differences in the reconstruction accuracy for the different features, we ran the regression model on the test set, and for each numerical feature, we computed the mean absolute error. To simplify the comparison, we used the normalized features and averaged together the errors for different features belonging to the same examination (e.g., the various questions in the ADAS questionnaire). The results are reported in Figure 2b and clearly show that the mean absolute error varies significantly between the different examinations. This difference is primarily due to some examinations having more samples available in the dataset; indeed, a linear model which uses samples^-1^ as a regressor to predict the mean absolute error has an *R*^2^ = 0.42. The mean absolute error (MAE) for any given numerical feature can be decomposed into two parts:

- Initial Estimation Error (IEE), which is the error due to the estimation of the patient state at the first time step (i.e., the first visit of the patient).
- Evolution Error (EE), which is the error due to the evolution of the patient’s state over time.

We can distinguish these two parts of the error fitting a linear model MAE = IEE + EE Δ, where Δ is the number of years between the first visit and the visit for which we are computing the error. In Figure 2c, we report the values of IEE and EE for several features, which we estimated by fitting the linear model on the predictions obtained using the network on the test set. We notice that for some features, the dominant error is either the IEE or the EE components, which is consistent with the different medical meanings of the features. For example, the APOE4 feature is a genetic feature that does not change over time; thus, the IEE component is dominant. On the contrary, the FDG features quantify the functional damage to the brain, so they evolve in hard-to-predict ways; thus, the EE component is dominant.

### Typical trajectories

To understand which temporal patterns the model has learned, we plot the typical trends of each biomarker for healthy, MCI, and dementia patients, cf.

Figure 3a. For each group, we show the mean predicted trajectories of a selection of features and their respective standard deviations. For some features, the trends are significantly different between the three groups (e.g., MMSE and CDRSB), while for others, the trends are closer (e.g., cortical volume increase). The trajectories of the different features are not independent. To visualize the relationships that the model has learned, we can plot the average vector field in the space determined by two features. To do this, we computed the derivatives of the predicted trajectories of each patient in the test set at the time of each visit, then for each pair of variables, we subdivided the corresponding plane into a grid and we averaged together the derivatives corresponding to predicted values that lay in the same cell of the grid. In Figure 4 we show the stream plots of a few features computed with this approach; in particular, looking at the plot in the space of amyloid-PET and tau-PET levels, we notice that the model has learned the typical amyloid cascade pattern (an increased amyloid levels is followed by an increase in tau levels).

**Figure 3.**
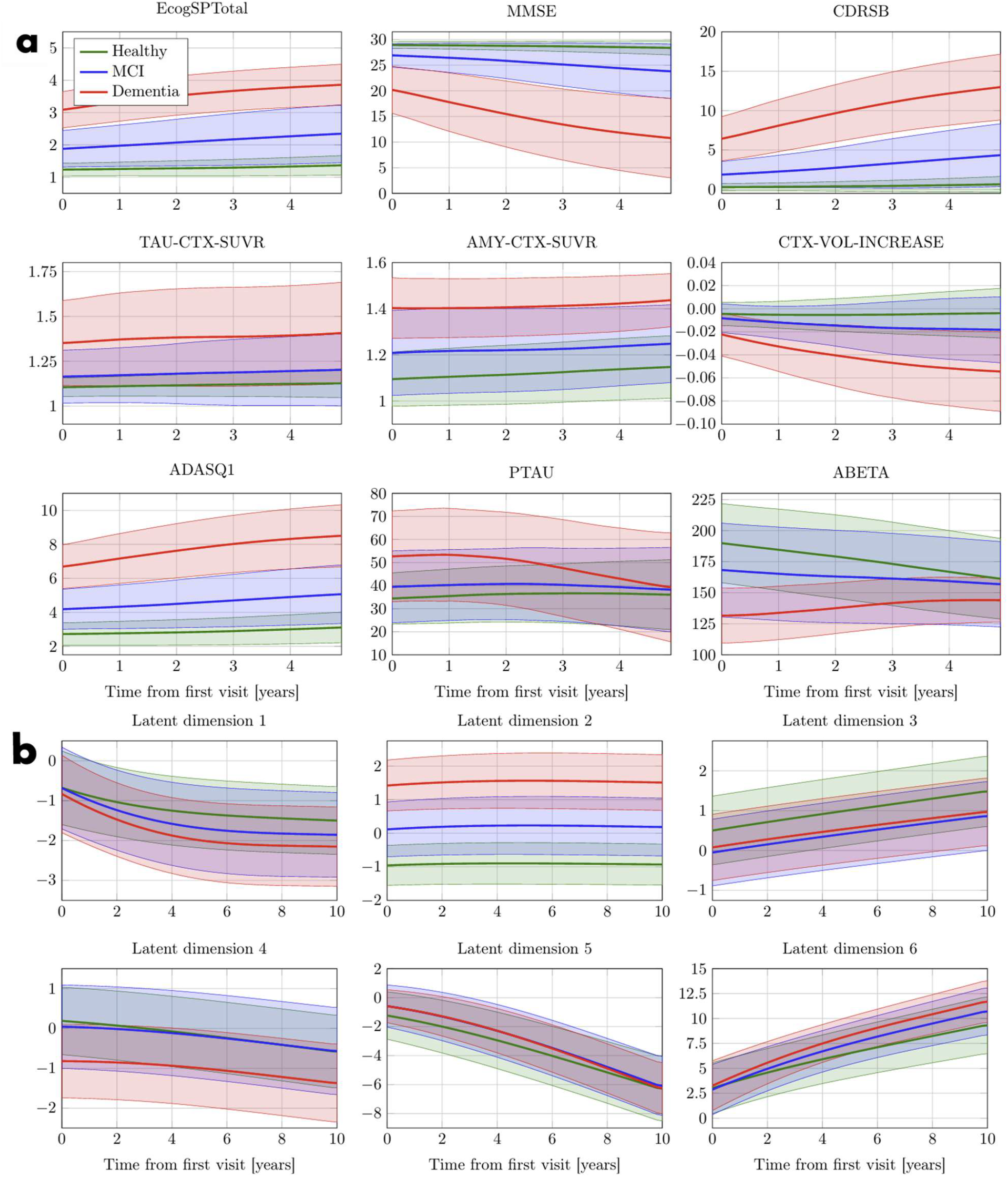
Average trajectories for healthy, MCI, and dementia patients. The lines represent the means of the predicted time series, and the shaded areas represent the standard deviation of the predicted time series. **a**, typical trajectories of selected AD markers. **b**, Average trajectories of the latent state variables. Refer to Table 4 for the list of acronyms.

**Figure 4.**
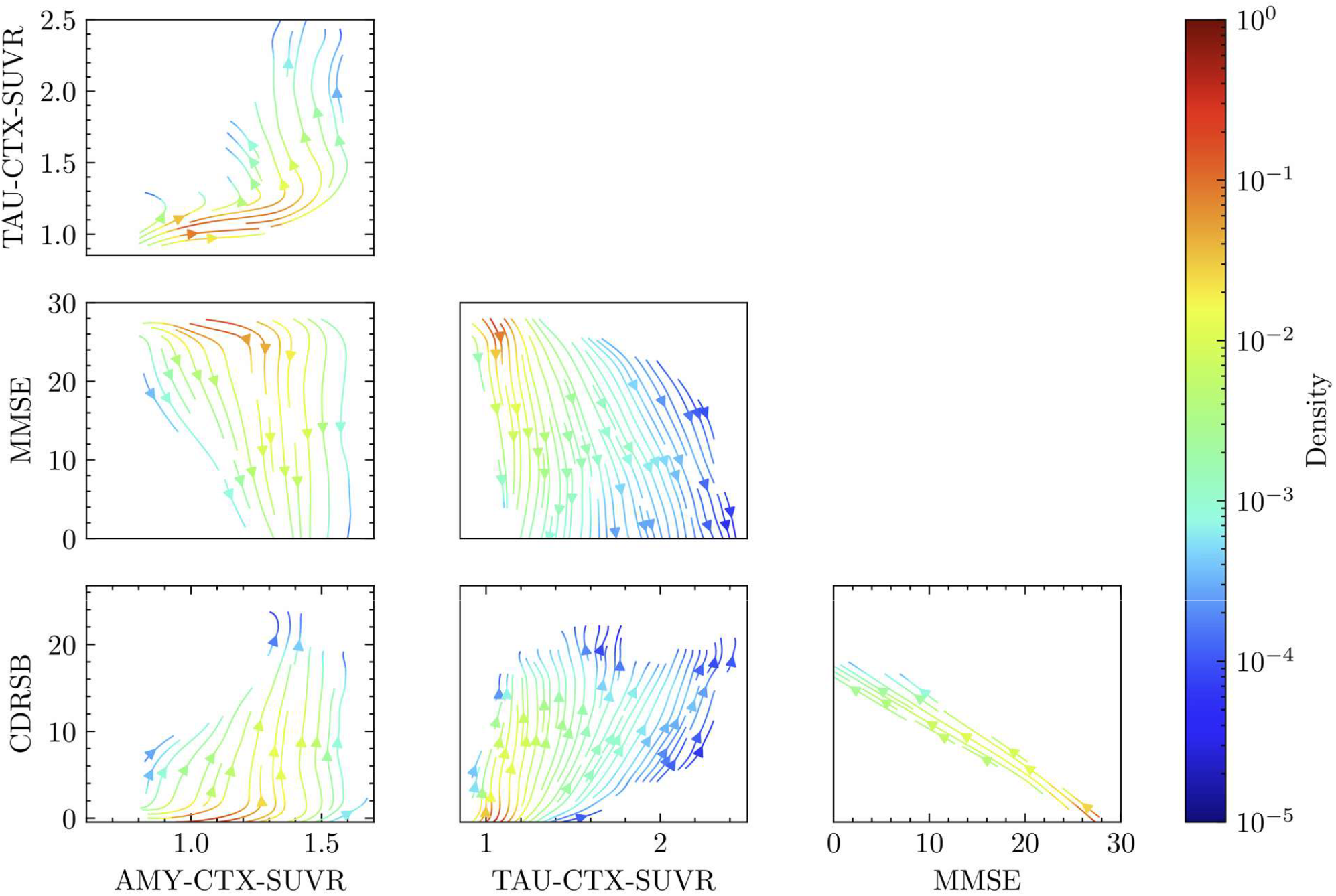
Stream plot of the average evolution of the amyloid-PET, tau-PET, MMSE, and CDR scores. The streamlines fill the space of values for which there are samples in the dataset, and the color represents the density of the samples. Refer to Table 4 for the list of acronyms.

### Latent space analysis

In our framework, the latent vector **z**(*t*) ∈ ℝ^*d*^ represents a compact numerical representation from which the DECODER-NN can reconstruct the multimodal panel of examination at time *t* (**x**(*t*)). Since the training is unsupervised and end-to-end, the model is forced to generate different latent trajectories between patients characterized by different observable AD marker evolutions. Since this operation is done in a compact space (*d* is usually selected small enough to avoid overfitting), the latent variable provides a new point of view on the hidden course of AD progression beyond the single disease timeline. This automatically opens to subtype differentiation and also benefit early diagnosis. Since a direct clinical interpretation is not possible, we report in this section the post-processing techniques used to showcase the relationship between **z**(*t*) and **x**(*t*).

Our model represents the patient state as a vector in a 6-dimensional latent space. This setting allowed for the best balance between accuracy and interpretability (see the Methods section for further details). We remark that using too few latent variables might underestimate the complexity of AD progression (under-fitting) and fail to capture the interplay of the distinct biological processes measured through the observable biomarkers. Conversely, a too large latent space can lead to overfitting and reduced interpretability. To investigate the trends characterizing these latent variables, we plotted the average evolution of each variable for healthy, MCI, and dementia patients (Figure *3*b). To get a more straightforward interpretation of what each variable represents, we used JAX’s automatic differentiation to compute the average Jacobian matrices of the normalized features reconstructed at the current time and after 5 years concerning the normalized latent variables at the current time (Figure 5). The two Jacobian matrices show us how sensitive each feature is to changes in the latent vector elements

**Figure 5.**
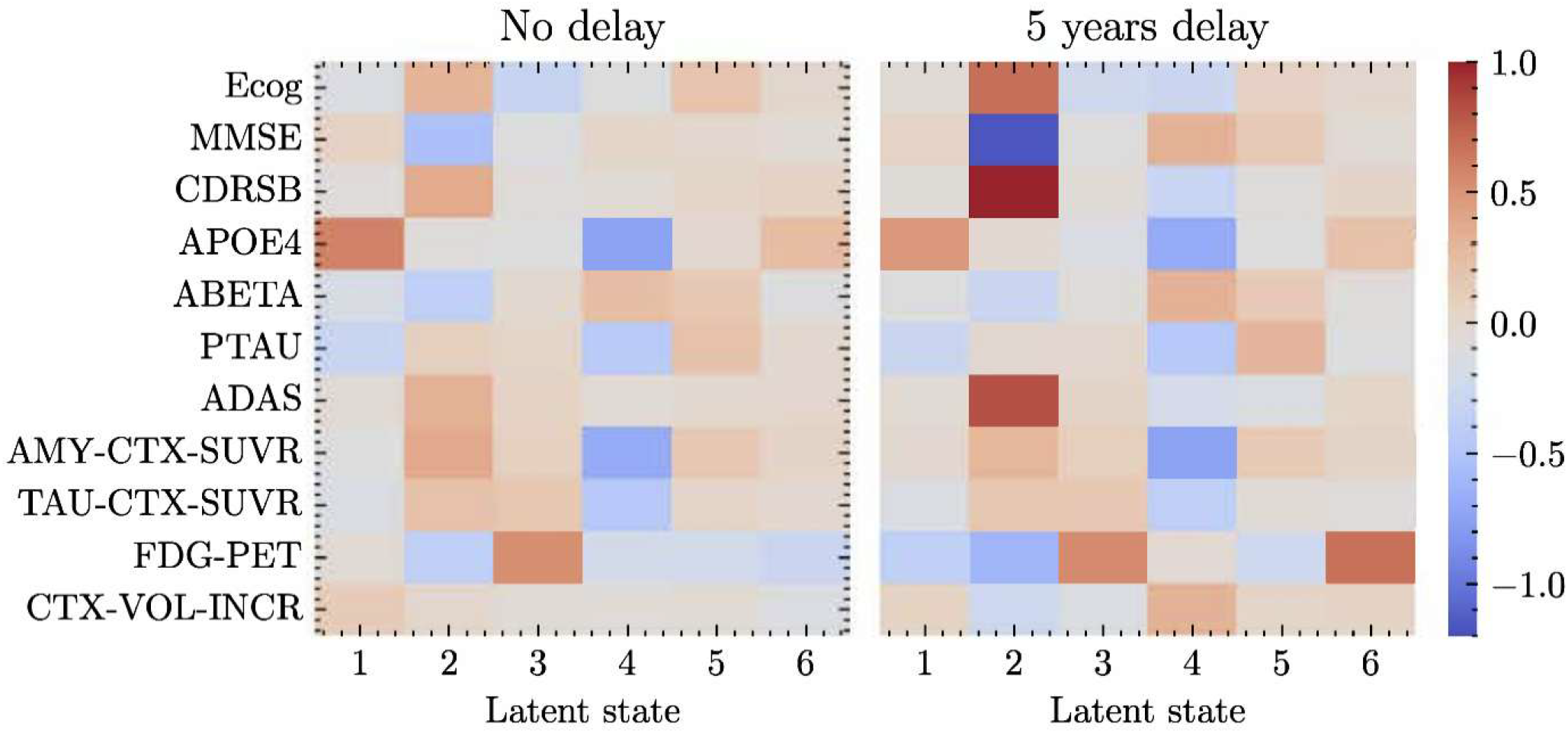
Average Jacobian matrices of several normalized AD markers reconstructed at the current time and after 5 years w.r.t. the normalized latent variables at the current time. The features belonging to the same exam (e.g., the questions of the ADAS questionnaire) have been averaged together. Refer to Table 4 for the meaning of the acronyms.

### Confidence Bands

To provide interpretable results for the predicted time series, we set a simultaneous confidence level so that there is at least an 80% chance that the entire series lies within the band. These bands are wider than pointwise intervals but easier to communicate. Since our model is generative, we can sample multiple future trajectories for each patient to estimate the mean and variance at each time step. We use these statistics and an empirical scaling factor derived from the validation set to determine confidence intervals that cover the whole time series. This approach allows us to assess confidence bands that do not depend on the number of predicted time steps and naturally account for correlations between time points, unlike simpler methods such as the Bonferroni correction. Figure 6 shows the confidence bands for the predicted time series of three patients from the test set, one for each possible outcome: healthy, MCI, and dementia.

**Figure 6.**
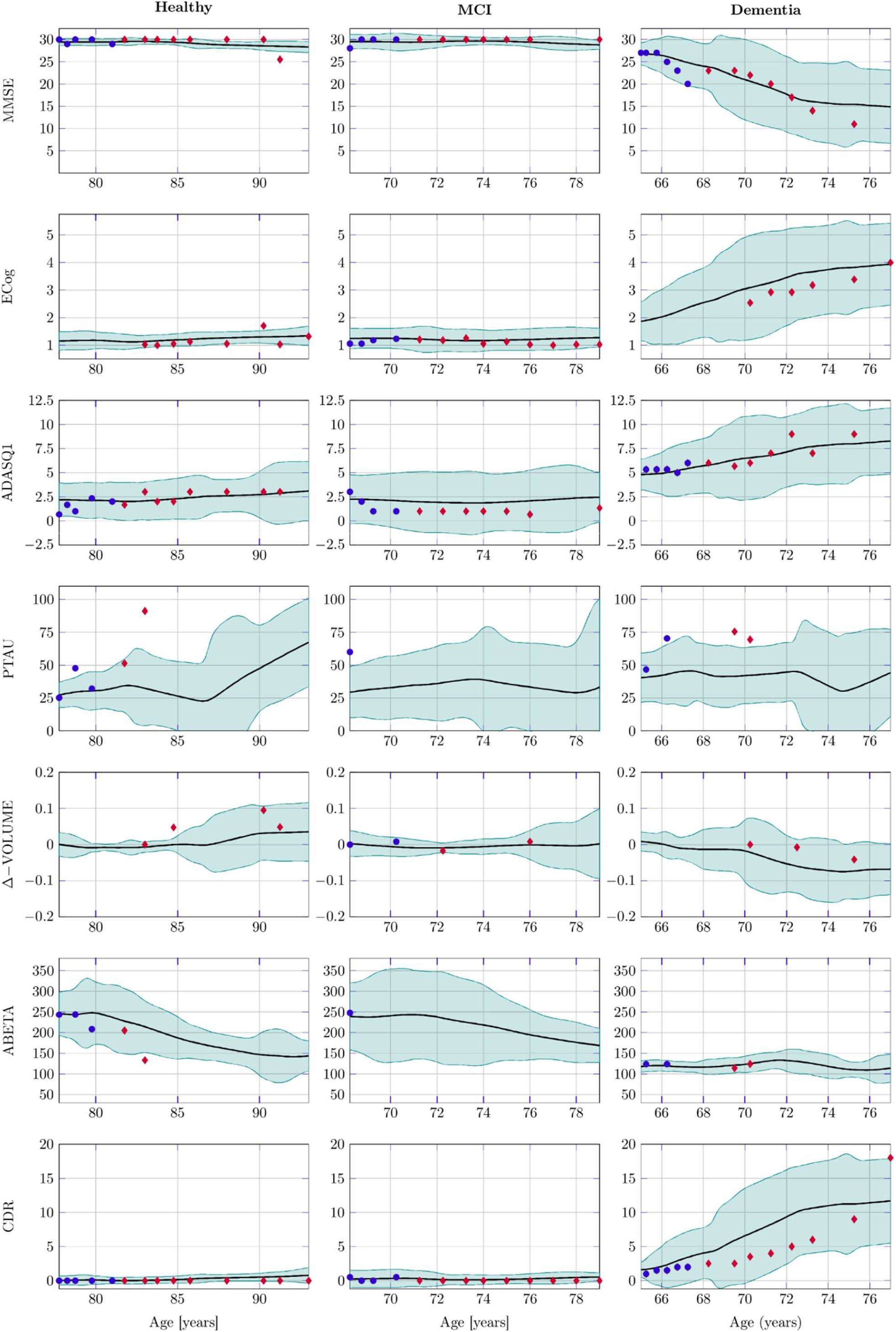
Prediction of several features for three patients: healthy (left), MCI (center), and dementia (right). The line represents the mean of the predicted time series with the corresponding confidence band, the circles represent the input values, and the diamonds represent the future values. Refer to Table 4 for the meaning of the acronyms.

## Discussion

In this work, we have introduced a new data-driven Disease Prediction Model for Alzheimer’s Disease based on NeuralODE architectures and the techniques to train it. The encoder, based on the ODE-RNN architecture, first identifies a suitable patient-specific initial latent state or its distribution at the time of the first visit. This design choice aligns with the fact that patients presenting at the first examinations can be at different stages of disease progression and removes the need for warping functions and time shifts in the model. Instead of displaying a one-dimensional disease timeline, our model identifies a continuous latent vector **z**(*t*) ∈ ℝ^*d*^, creating a reduced-order manifold of disease evolution (see Figure 3b and Figure 4). This enables us to capture nonlinear relationships between heterogeneous features and identify different trajectories in the same patient’ group, that are not limited to how fast the disease progresses or the stage at which the patient presents. Finally, the decoder is responsible for the time integration of the latent vector starting from the initial condition provided by the encoder and the reconstruction of the biomarkers’ trajectories or the classification of the patients.

The model comparison reported in Table 3 (Methods) shows that the NeuralODE-based DPM models significantly outperform the LSTM model in all the metrics; this result confirms that for AD modelling, using NeuralODEs as the fundamental building block of the model is a better choice to handle the typical sampling heterogeneity of clinical data. Furthermore, the generative approach with a distribution estimation and sampling does not significantly impact the model performance, but it enables us to model and communicate the uncertainty of the predictions. This is an architectural improvement that can be integrated into other DPMs based on neural networks, such as the MC-RVAE [18] because the NeuralODE can act as a drop-in replacement for the LSTM layer or other recurrent neural networks. Regarding the dependency on the dataset size, the error rate plotted versus the number of patients in the training set reaches a plateau when using about 75 % of the available data. Therefore, the model complexity is adequate for the current dataset size.

The 74% precision and 52% recall achieved in the early diagnosis task are significant results. Indeed, the natural choice of using the current diagnosis as an estimate of the future diagnosis has a 67% precision and 0% recall. This result indicates that the model has captured some early risk factors that were not accounted for in the diagnosis process. There is still room for improvement in performing an early prediction of dementia, which is misclassified as MCI in most of the patients, but this result is expected since we are considering only data from the earliest stage of the disease.

The patterns learned by the model match the medical knowledge: for example, the MMSE score is stable for healthy subjects, decreases for MCI patients, and decreases even more rapidly for dementia patients (see Figure *3*a). Concerning MMSE, values of the trajectories resulting from our model are coherent with the literature values from medical examinations [22]. Trends coherent with medical knowledge were found also for other observable features, except for the CSF levels of Aβ and phosphorylated tau. They correctly present distinct trajectories at the beginning of the time window, but over extended time periods seem to converge to similar mean values. This behavior is consistent with the fact that, for the majority of patients, these biomarkers are typically assessed only once at baseline. For this reason, the training dataset is missing the longitudinal information needed for a data-driven model, which therefore provides the most parsimonious trajectory (the average behavior).

Concerning the interpretation of latent variables, we can notice the following relationships from the sensitivity analysis of Figure 5. The APOE4 gene is encoded mainly through the first and fourth latent variables. These sensitivities are also found when considering the 5-year delay in the output, matching the genetic nature of the APOE4 gene, which does not change over time. The latent variable two is strongly correlated with cognitive tests’ results, especially in the long term: variations of this variable are associated with an increase in ADAS, CDR, and ECog scores, and a decrease in MMSE scores, which indicate a worsening of the cognitive abilities. This latent variable is by far the one that shows the most significant differentiation between the three groups, as visible in *Figure 3*b. For this variable, the between-group variance [23] accounts for 46% of the total variance; on the contrary, in all the other variables, it accounts for less than 15%, although the cognitive tests tend to significantly worsen over time in MCI and dementia patients (Figure 3a). However, the latent variable 2 is stable over time; therefore, we can interpret it as a measure of the severity of the underlying disease, rather than a measure of cognitive impairment, which seems more associated with the first latent variable. The latent variables 3 and 4 are mainly linked to imaging and CSF biomarkers. Specifically, latent variable 3 captures the FDG PET scores. The fact that the model has learned to represent FDG PET in a separate latent variable suggests it may serve as an independent biomarker for the disease, in addition to the established ATN framework, which aligns with several previous studies [24–26]. Latent variable 4 encodes Aβ in the brain and CSF, total tau levels in the brain, and phosphorylated tau levels in the CSF. Notably, CSF Aβ and PET Aβ are affected in opposite ways, consistent with the underlying biological mechanisms. These features are also partially represented in latent variable 2. However, in Figure 3b, we see a separation of the dementia group from the other groups in latent variable 4, indicating that the model has learned Aβ and tau levels worsen more than cognitive abilities during the progression from MCI to dementia. Finally, the latent variables 5 and 6 change significantly over time, do not show a clear separation between the groups, and do not have a clear pattern of correlated features, so we may suppose that they probably encode for the concomitant ageing process. They are also by far the most correlated latent variables: their correlation is -0.82, while the other latent variables show at most a correlation of -0.36.

Analyzing the phase space plots in Figure 4, we can notice that the cortical Aβ accumulation (measured by the AMY-CTX-SUVR) increases before the tau protein accumulation, and the variation of MMSE and CDR. These patterns are coherent with the amyloid cascade hypothesis [4,27] and previous studies [12]. On the contrary, there is less delay between the tau protein accumulation and the change in the clinical test performances, and there is a strong correlation between the CDRSB and MMSE variation times. Also, this fact is coherent with the clinical development of Alzheimer’s disease [28].

Most DPMs based on neural networks do not provide a measure of the uncertainty in their predictions, which in many cases is quite high due to the small number of observations per patient and the high variability in disease manifestations. By sampling the latent state embedded in the model and using the approach presented in ‘Confidence bands’ section, our model can generate a confidence band for its predictions that is easily interpretable by doctors and communicable to patients. In ‘Model performance’ section, we also show that incorporating this uncertainty quantification does not reduce the average model’s accuracy. We designed the model architecture to be as flexible as possible across different tasks; for example, it has been used to reconstruct the trajectories of AD markers and to provide early diagnosis (see ‘Early diagnosis’ section). This flexibility, combined with its ability to handle large percentages of missing data, makes the architecture suitable as a basis for a foundational model that can be trained on large, unlabeled datasets with sparse observations.

We have shown that our model can successfully be trained to predict the evolution of 46 different features; typically, DPMs are developed on a much smaller number of examinations, and their structure is specifically designed for that set of examinations. Being able to handle several features and being completely data-driven make our model suitable as a research tool to investigate the predictive power of candidate biomarkers for AD. We have also shown a couple of techniques to extract insights from the model, which can guide future research on an ATX(N) system development [29]. The ability to be trained or fine-tuned on sparse datasets is a significant advantage, as economic and practical constraints often limit the number of enrolled patients and the examinations performed. As evidence of the model’s ability to identify functional patterns, in the ‘Typical trajectories’ section, we have shown that the model has autonomously learned the amyloid cascade dynamics [4] (decrease in cerebrospinal fluid Aβ followed by an increase in tau levels) without any prior knowledge or pre-imposed structure. This strongly indicates that the model can learn complex dynamics directly from data without human supervision. Lastly, we have demonstrated promising results in early diagnosis, which suggests that the model could be a valuable tool for improving early differential diagnosis of AD.

## Methods

In this section, we detail our approach to developing a disease progression model for clinical applications. Specifically, we describe the structure of our model, the reasons behind our architectural choices, the implementation details, dataset preprocessing, and the training process. To tackle these challenges, we adopted a novel approach based on NeuralODEs, which, to the best of our knowledge, has not been previously used in the context of neurodegenerative diseases.

### Model architecture

For any given patient, the input data that the model receives is the sequence of the medical examinations the patient has had so far, indexed by the patient’s ages *t*_0_, …, *t*_*n*_ at the time of each visit. The model we developed represents the patient state at time *t* as a latent vector **z**(*t*) ∈ ℝ^*d*^. Representing the patient state as a low-dimensional latent variable, instead of using the observed visits directly, allows the model to be trained on high-dimensional data without overfitting because the low-dimensional latent state acts similarly to the bottleneck layer in an autoencoder. The latent state is evolved using a NeuralODE, which is an ODE whose right-hand side is defined by a neural network 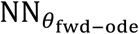 (in this section the vectors *θ* represent the weights and biases of the corresponding neural network). The NeuralODE layer is implemented using a numerical solver to approximate the ODE problem using Tsitouras’ 5/4 method [30], which is a Runge-Kutta method with adaptive step size. The model includes a fully connected neural network 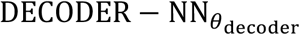 which maps **z**(*t*) to **x**(*t*), that is, it predicts the features’ vector from the latent state.

Finally, the model uses an encoder architecture (Algorithm 1) inspired to the ODE-RNN introduced in [19] to estimate the probability distribution of the latent state at the time of the first visit; this architecture allows the model to account for all the information from the visits the patient has had up to the present time (Figure 1), even though the exams are irregular in number and spacing. The encoder receives as input all the visits up to the present time and outputs the vector **Z** ∈ ℝ^2*d*^, which is interpreted as the parameters of a distribution over the latent states: the distribution is a multivariate Gaussian with a diagonal covariance matrix, and the vector **Z** is a concatenation of the mean vector and the log-variance vector. Estimating a distribution instead of a single latent state allows the network to be generative and model the inherent uncertainty in the prediction. A fully connected network 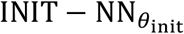 is initialized using as input the patient’s age at the last visit. Each visit is processed going backwards in time: a fully-connected neural network 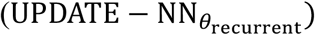 updates the latent state using the visit data and the current state, and between visits, the state is evolved using a NeuralODE 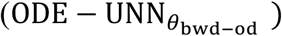 This architecture is similar to a recurrent neural network, but the hidden state evolves continuously, instead of being fixed between visits.

The examination data **x** is used as input only for 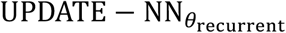 and the rest of the model works only with the latent state, so handling the missing values is trivial: all the NA values are replaced with zeroes and 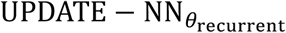 learns to encode the sparse visit vector as a dense latent state.

#### Algorithm 1: The ENCODER layer, which is our adaptation of the ODE-RNN algorithm proposed in [19].

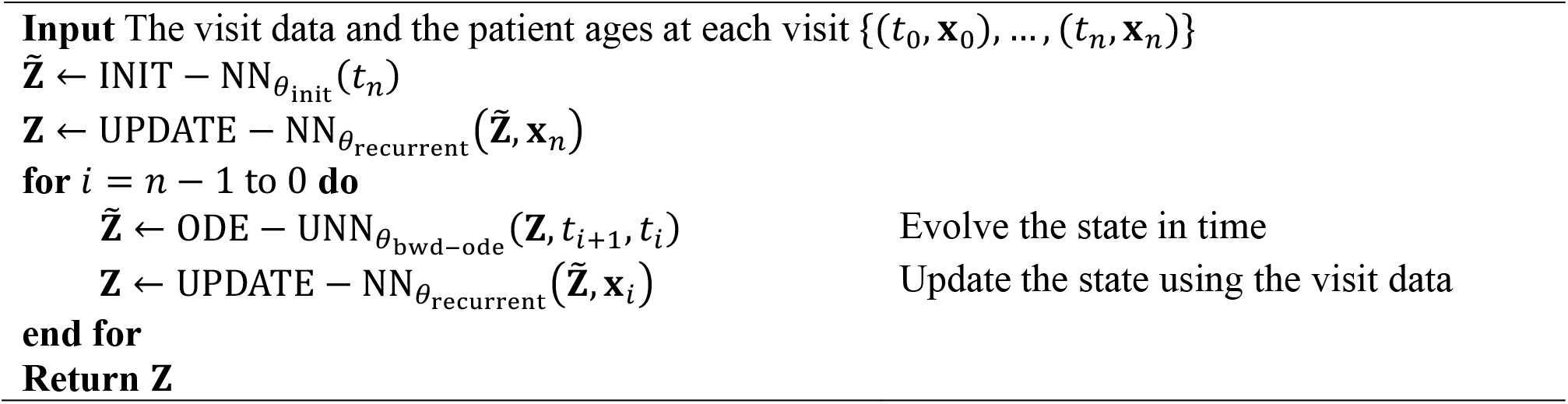

The complete model architecture, represented in Figure 1, is the following: the ODE-RNN layer encodes all the visits as a distribution over the latent state, the latent state is sampled from this distribution, the NeuralODE layer evolves the latent state in time and the decoder predicts the examination vectors at each time step of interest; which can be formalized as:

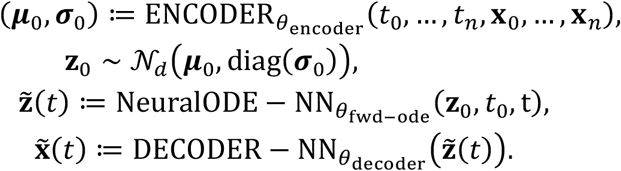

Here 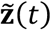 and 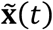are the predicted latent state and the visit data as functions of the patient’s age. We call this architecture Ordinary Differential Equation Autoencoder (ODE-AE), respectively.

### Dataset and data preprocessing

Data used in the preparation of this article were obtained from the Alzheimer’s Disease Neuroimaging Initiative (ADNI) database (adni.loni.usc.edu). The ADNI was launched in 2003 as a public-private partnership, led by Principal Investigator Michael W. Weiner, MD. The primary goal of ADNI has been to test whether serial magnetic resonance imaging (MRI), positron emission tomography (PET), other biological markers, and clinical and neuropsychological assessment can be combined to measure the progression of mild cognitive impairment (MCI) and early Alzheimer’s disease (AD).

The model was trained on the data from all the ADNI cohorts available at the time of training: ADNI1, ADNI GO, ADNI2, and ADNI3. The different cohorts followed different exam schedules and some features are available only for specific cohorts, e.g., the Neuropsychiatric Inventory assessment is available only for ADNI2 and ADNI3 patients. The total number of visits, the distance between visits, and the features available at each visit are patient-specific. We divided the patients into three groups: training (2451 patients), validation (613 patients), and test (766 patients). Then, we generated all the truncated time series for each patient. Splitting the patients instead of the time series ensures that data from the same patient does not appear in both training and test sets, preventing biased results. The results shown in the dedicated section were obtained using the test set to evaluate the model and the validation set to tune hyperparameters, such as the architecture’s dimensions and the weights of various losses.

We selected a subset of the dataset features representative of the brain’s physiological, structural, and functional status; these include demographic information, the diagnosis, neuropsychological assessments, neuroimaging data, genetic data, and cerebrospinal fluid biomarkers. For the complete list of features, we refer to Table 1 and the ‘Dataset features and acronyms’ section. The Tau PET and Amyloid PET data were preprocessed to calculate the standardized uptake value ratio (SUVR) over the cortical region, while the parietal and temporal SUVR were derived from the FDG PET data. The MRI data were used to determine the percentage change in hippocampal and cortical volumes compared to the initial measurement for each patient. To reduce the percentage of missing data (and thus speed up the training process) we aggregated all the visits of the same patient in the same trimester of the year into a single sample containing all the examinations performed in that trimester; examinations performed more than once in the same trimester were aggregated using the mean for numerical features and majority voting for categorical features. We replicated all the features that do not change over time (e.g., gender, genetic data, …) to all the visits of the same patient. Furthermore, all the data has been normalized before training the model.

### Training

We implemented the model using the JAX framework [31], which allowed us to handle the variable-length time series in input and output and to train the model end-to-end by backpropagating through the ODE numerical solver. The training process is unsupervised: the model learns to reconstruct all the visit data at all the time points using as input some of the visits up to a cutoff time. Given that the available visit data is sparse and different features have different percentages of missing values, we implemented a masking of the loss function: the error of each feature is averaged only over the subset of samples that contain that feature. Using this masked average approach ensures that all features have the same weight in the loss function, even if the percentage of missing values is different. In the following definitions, we use this notation:

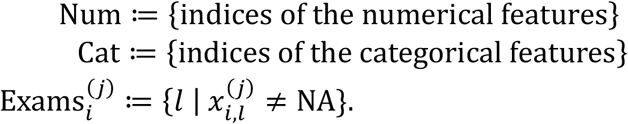

In particular, the indicator function 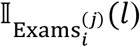 is equal to 1 if and only if the feature *l* is present in the *i*-th visit of patient *j*. The terms of the loss function used to train the model are the mean squared error (MSE) on each numerical feature

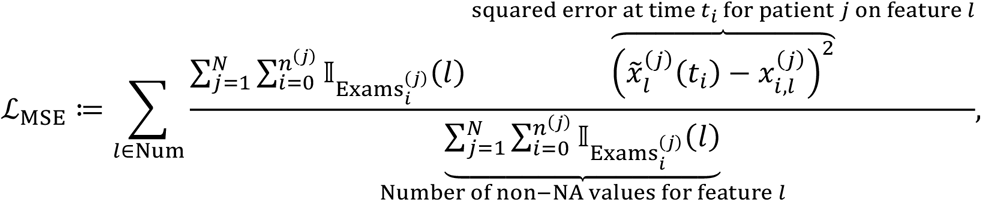

the categorical cross-entropy (CCE) on each categorical feature

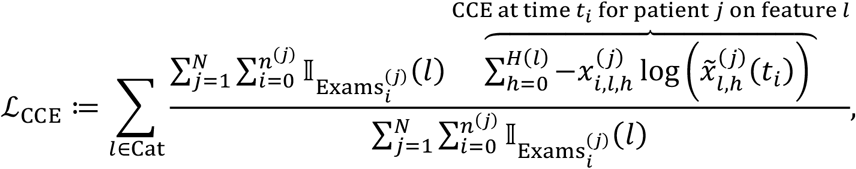

the *L*_2_ regularization on the NeuralODE’s weights

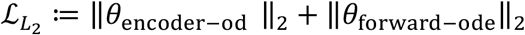

and the Kullback-Leibler (KL) divergence between the predicted latent state distribution and the standard Gaussian distribution

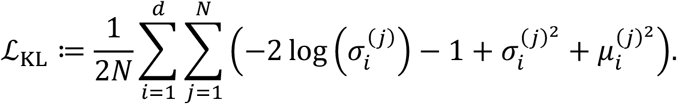

The loss function is defined as a weighted sum of all the terms:

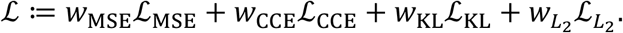

The MSE and CCE account for the reconstruction error, while the KL divergence is used to ensure the estimated distribution does not degenerate to a singular distribution; this is conceptually similar to the evidence lower bound (ELBO) used as loss in variational autoencoders [32]. Finally, the *L*_2_ regularization is used to prevent overfitting and to ensure the NeuralODE reconstructs parsimonious and regular dynamics in the latent space, which also makes numerical integration easier.

The training process used truncated time series 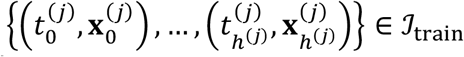 as input and the full time series 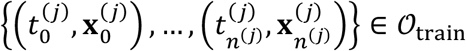 as target; this approach forces the model to learn to interpolate between observed visits and to predict future visits. The training set was obtained by selecting a random subset of patients and truncating their time series at each possible visit. Then, a random masking was applied to the input data to ensure the model learns to estimate also the missing features.

To parallelize the training process, even though the time series have different lengths, we padded the input data with NA values up to the maximum time series length; the model was implemented ignoring the time steps with NA timestamps both in input and output. This approach allowed us to parallelize the computation efficiently; however, it increased the memory requirements. Due to the limited size of the dataset, this trade-off was favorable. The backpropagation is more complex due to the presence of the NeuralODEs and the random sampling step.

Since the random variables follow Gaussian distributions, we can use the reparameterization trick by [32] to fall back into the standard back-propagation setting: represent the random variable as 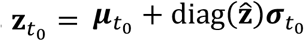 with 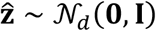, then the random variable 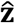 does not depend on the model parameters and as such it can be considered constant when computing the gradient for the gradient descent step.

To fit the NeuralODEs, we used diffrax to implement a discretize-then-optimize approach [33]: we integrate the ODE numerically as part of the model implementation and then backpropagate through the numerical integration steps using JAX’s automatic differentiation features. This was the optimal choice for our use case because it is accurate and fast, with the only drawback being memory inefficiency, which is not a concern in our case since the time horizon is short, and the number of time steps of interest is low. These techniques allowed us to train the model in an end-to-end fashion, without the need to split it into multiple parts and use different optimization techniques for each part. We tuned the hyperparameters of the model using a grid search approach; in Table 2, we report some of the hyperparameters we tested and the resulting model performance on the validation set. We notice that the accuracy of several models is very similar; for all the manuscript analyses, we used the model with six latent dimensions and a single 16-neuron layer in the NeuralODE (the first one in Table 2), because it had the best performance and was the most parsimonious. To ensure that the training process was robust, we retrained the model 5 times with different random initializations. All the models had reconstruction errors in the [11.6,12.1] range on the validation set, which confirms the stability of the training process. As part of the hyperparameter tuning, we tested three different optimizers: Adam [34], L-BFGS [35], and RMSProp; the Adam optimizer with an exponential learning rate decay was the best performing one, and all the reported experiments were performed using it.

**Table 2.** Reconstruction errors of the model trained with different hyperparameters. The errors were computed on the validation set using the ME approach (‘Evaluation’ section).

|  |  |  |  |  |  |
| --- | --- | --- | --- | --- | --- |
| **Number of Latent Dimensions** | 6 | 6 | 12 | 6 | 6 |
| **NeuralODE neurons** | 16 | 24,24 | 24,24 | 16 | 16 |
| **$L_2$ regularization weight** | 0.02 | 0.02 | 0.2 | 0.02 | 0.02 |
| **KL divergence weight** | 1 | 1 | 1 | 10 | 1 |
| **Initial learning rate** | 0.003 | 0.003 | 0.003 | 0.003 | 0.01 |
| **Learning rate decay** | 0.97 | 0.97 | 0.97 | 0.97 | 0.97 |
| **Learning rate decay steps** | 2000 | 2000 | 2000 | 2000 | 2000 |
| **Reconstruction error** | 11.64 | 11.71 | 11.65 | 13.05 | 13.06 |

### Evaluation

The model estimates a Gaussian distribution 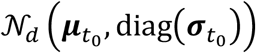 over the latent states, which defines a distribution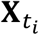 over the visit data at any time *t*_*i*_. At inference time we are interested in the mean of 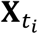 which we estimated in two ways:

- Monte Carlo (MC): *B* latent states are sampled from the distribution, and for each latent state, the model predicts the visit data; the final prediction is the mean of the *B* predictions. We can formalize this as:

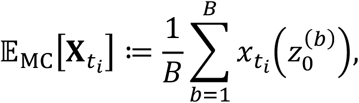

where 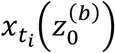 is the predicted visit data at time *t*_*i*_ obtained using the *b*-th sampled latent state.
- Mean estimation (ME) :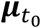 is used as the initial latent state and the visit data is predicted deterministically from this mean latent state, which can be formalized as

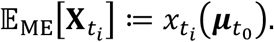

In principle, the two approaches could produce different results because the mean latent state does not necessarily match the mean predicted visit data. However, if the transformation from the latent state to the visit data is smooth with small second derivative then it should be close. We expect this assumption to hold because of the regularization used during training, and in practice, we found that the MC and ME approaches yield very similar results. Therefore, we chose the ME approach since it is more computationally efficient.

We used as evaluation metric the reconstruction error, which includes the MSE and categorical cross-entropy losses, but does not include the KL divergence and the *L*_2_ regularization; we computed the reconstruction error on the test set using the ME approach. To evaluate the overall model performance, we computed the reconstruction error on the whole time series, including the input data, while to assess the forecasting performance, we used only the future visits data, which the model didn’t receive as input.

### Model comparison

We compare our new model to an alternative based on the Long Short-Term Memory (LSTM) architecture [36], which is one of the most commonly used recurrent neural network architectures to model time series and has been used in several works on AD progression modelling [37–39]. To evaluate the performance of the ODE-AE architecture, we compared it with two alternatives:

- **LSTM**: a model that uses LSTM layers [36] instead of the NeuralODE and the ODE-RNN layers.
- **Deterministic version**: a simplified version of ODE-AE which does not include the distribution estimation and the sampling step, but still uses the NeuralODE and the ODE-RNN layers.

We define the reconstruction error as the combined sum of the MSE for the numerical features and CCE for the categorical features (see ‘Training’ section). Because the models can interpolate the patient state up to the present and forecast future states, we evaluated them using three different metrics: the interpolation error, prediction error, and overall error. The interpolation error is the reconstruction error computed on the visits up to the present. The prediction error is the reconstruction error computed on the future visits. The overall error considers all the visits. The results of the evaluation are reported in Table 3.

**Table 3.**
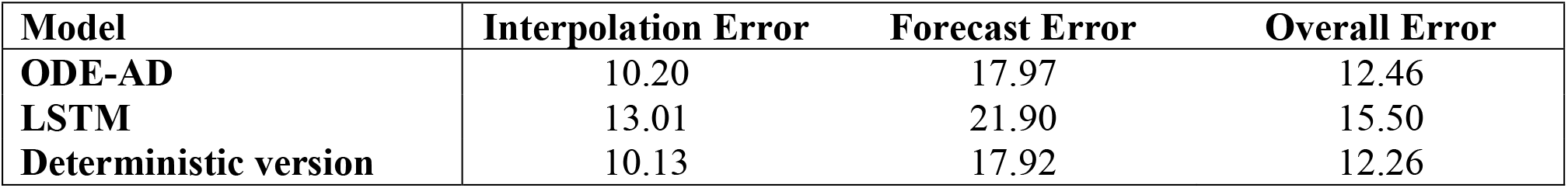
Reconstruction errors of the different models on the test set.

| Model | Interpolation Error | Forecast Error | Overall Error |
| --- | --- | --- | --- |
| **ODE-AD** | 10.20 | 17.97 | 12.46 |
| **LSTM** | 13.01 | 21.90 | 15.50 |
| **Deterministic version** | 10.13 | 17.92 | 12.26 |

Furthermore, to assess the effect of the number of training patients on the model performance, we trained the model 6 times with different numbers of patients, and for each model, we computed the interpolation error and the forecast error on the test set; the results are shown in Figure 7. Interpolation and forecast errors versus the number of patients in the training set..

**Figure 7.**
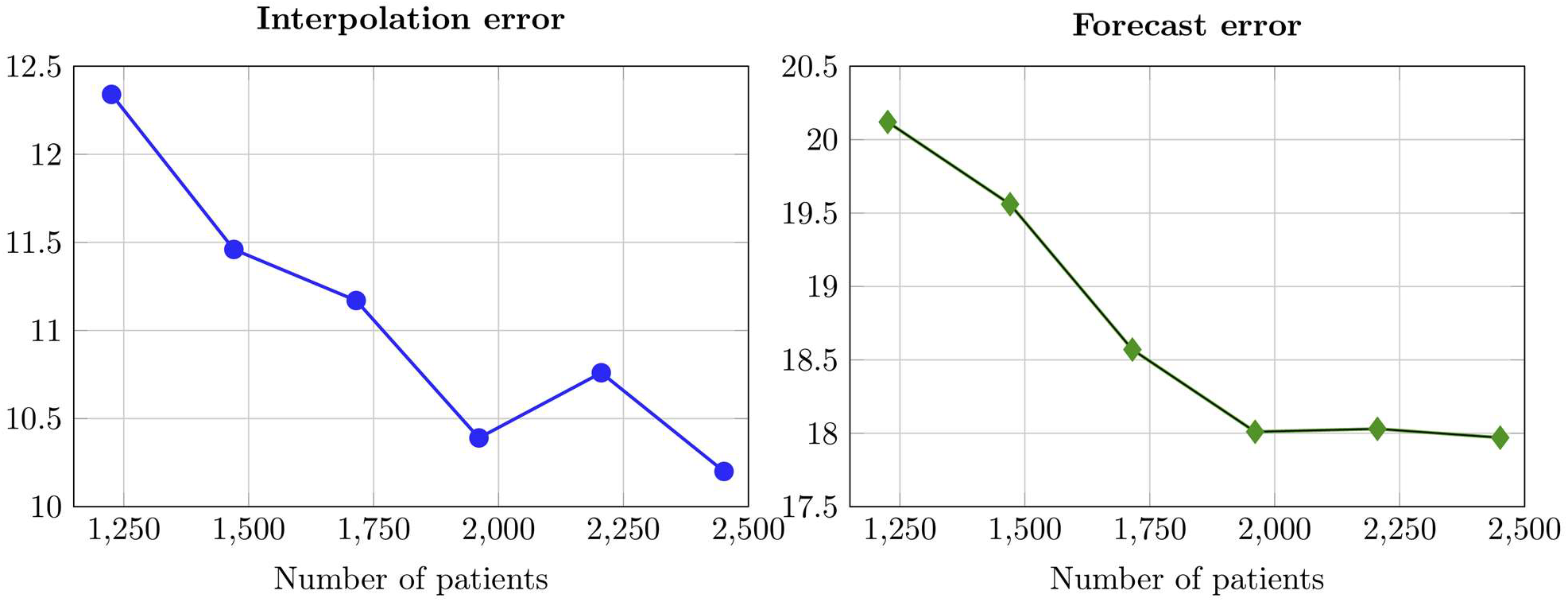
Interpolation and forecast errors versus the number of patients in the training set.

### Dataset features and acronyms

The dataset extracted from the ADNI database contains 3,830 patients with a total of 15,851 visits (after aggregation by trimester). The dataset includes the following features:

- Demographic: gender, age, marital status, years of education, handedness.
- Diagnosis: a categorical variable with values: healthy, MCI, dementia.
- Neuropsychological assessments: Mini-Mental State Examination (MMSE) total score, Clinical Dementia Rating (CDR) total score, Everyday Cognition (ECog) total score differentiated between self-administered and study partner-administered tests, Neuropsychiatric Inventory (NPI) total score, Auditory verbal Rey test score, Alzheimer’s Disease Assessment Scale (ADAS) scores for each question, Clock Drawing total score, Digit span score for forward and backward test, Category fluency score in each sub-test.
- Neuroimaging data: Tau PET and Amyloid PET standardized uptake value ratio (SUVR) over the cortical region, FDG PET SUVR over the parietal and temporal regions, MRI percentage variation of the hippocampal and cortical volumes compared to the first measurement
- Genetic data: APOE4 gene presence.
- Cerebrospinal fluid (CSF) biomarkers: amyloid-β42, amyloid-β40, phosphorylated tau.

This wide range of features allows the model to learn the etiology and pathophysiology of the disease. Throughout this work, we use the acronyms listed in Table 4 to refer to the features of the dataset.

**Table 4.** List of acronyms used in the dataset.

| Acronym | Description |
| --- | --- |
| MMSE | Mini-Mental State Examination |
| CDR, CDRSB | Clinical Dementia Rating |
| ECog, EcogSPTotal | Everyday Cognition |
| ADAS | Alzheimer's Disease Assessment Scale |
| ADASQ1 | First question of the ADAS questionnaire |
| APOE4 | Apolipoprotein E4 |
| CSF | Cerebrospinal Fluid |
| CLOCKSCOR | Clock Drawing Test |
| CAT | Category Fluency Test |
| DSPAN | Digit Span Test |
| ABETA | Amyloid-β in the cerebrospinal fluid |
| PTAU | Phosphorylated Tau in the cerebrospinal fluid |
| SUVR | Standardized Uptake Value Ratio |
| MRI | Magnetic Resonance Imaging |
| PET | Positron Emission Tomography |
| AMY-CTX-SUVR | Amyloid PET SUVR over the cortical region |
| TAU-CTX-SUVR | Tau PET SUVR over the cortical region |
| FDG-PET | Fluorodeoxyglucose PET |
| FDG-PARIETAL | FDG PET SUVR over the parietal region |
| FDG-TEMPORAL | FDG PET SUVR over the temporal region |
| Δ-VOLUME,<br>CTX-VOL-INCREASE,<br>CTX-VOL-INCR | Increase of the cortical volume relative to the first measurement |
| HIPPO-VOL-INCR | Increase of the hippocampal volume relative to the first measurement |

## Acknowledgements

P.F.A. has been partially funded by ICSC—Centro Nazionale di Ricerca in High Performance Computing, Big Data, and Quantum Computing, funded by the European Union— NextGenerationEU. PFA, MC and SP are members of INdAM-GNCS. The present research is part of the activities of “Dipartimento di Eccellenza 2023-2027”, MUR, Italy.

**ADNI Acknowledgements**

Data used in the preparation of this article were obtained from the Alzheimer’s Disease Neuroimaging Initiative (ADNI) database (adni.loni.usc.edu). The ADNI was launched in 2003 as a public-private partnership, led by Principal Investigator Michael W. Weiner, MD. The primary goal of ADNI has been to test whether serial magnetic resonance imaging (MRI), positron emission tomography (PET), other biological markers, and clinical and neuropsychological assessment can be combined to measure the progression of mild cognitive impairment (MCI) and early Alzheimer’s disease (AD). For up-to-date information, see www.adni-info.org.

Data collection and sharing for this project was funded by the Alzheimer’s Disease Neuroimaging Initiative (ADNI) (National Institutes of Health Grant U01 AG024904) and DOD ADNI (Department of Defense award number W81XWH-12-2-0012). ADNI is funded by the National Institute on Aging, the National Institute of Biomedical Imaging and Bioengineering, and through generous contributions from the following: AbbVie, Alzheimer’s Association; Alzheimer’s Drug Discovery Foundation; Araclon Biotech; BioClinica, Inc.; Biogen; Bristol-Myers Squibb Company; CereSpir, Inc.; Eisai Inc.; Elan Pharmaceuticals, Inc.; Eli Lilly and Company; EuroImmun; F. Hoffmann-La Roche Ltd and its affiliated company Genentech, Inc.; Fujirebio; GE Healthcare; IXICO Ltd.; Janssen Alzheimer Immunotherapy Research & Development, LLC.; Johnson & Johnson Pharmaceutical Research & Development LLC.; Lumosity; Lundbeck; Merck & Co., Inc.; Meso Scale Diagnostics, LLC.; NeuroRx Research; Neurotrack Technologies; Novartis Pharmaceuticals Corporation; Pfizer Inc.; Piramal Imaging; Servier; Takeda Pharmaceutical Company; and Transition Therapeutics. The Canadian Institutes of Health Research is providing funds to support ADNI clinical sites in Canada. Private sector contributions are facilitated by the Foundation for the National Institutes of Health (www.fnih.org). The grantee organization is the Northern California Institute for Research and Education, and the study is coordinated by the Alzheimer’s Disease Cooperative Study at the University of California, San Diego. ADNI data are disseminated by the Laboratory for Neuro Imaging at the University of Southern California.

## Notes

### Competing Interest Statement

The authors have declared no competing interest.

